# Lipid metabolic perturbation is an early-onset phenotype in adult *spin* mutants: a *Drosophila* model for lysosomal storage disorders

**DOI:** 10.1101/075556

**Authors:** Sarita Hebbar, Avinash Khandelwal, R Jayashree, Samantha J. Hindle, Yin Ning Chiang, Joanne Y. Yew, Sean T. Sweeney, Dominik Schwudke

**Author notes:** Equal contribution as corresponding authors. Present Address Max Planck Institute for Molecular Cell Biology and Genetics, Dresden 01277 (Sarita Hebbar). Present Address EMBL/CRG Systems Biology Research Unit, Centre for Genomic Regulation (CRG), Dr. Aiguader 88, 08003 Barcelona, Spain & Universitat Pompeu Fabra (UPF), Barcelona, Spain. Present Address Research Center Borstel, Borstel, 23845, Germany (Dominik Schwudke). Department of Anesthesia and Perioperative Care, Genentech Hall, 600 16^th^ Street, University of California San Francisco, San Francisco, CA.

## Abstract

Intracellular accumulation of lipids and swollen dysfunctional lysosomes are linked to several neurodegenerative diseases including lysosomal storage disorders (LSD). A detailed characterization of lipid metabolic changes in relation to the onset and progression of neurodegeneration is currently missing. In this study, we systematically analyzed lipid perturbations in *spinster (spin)* mutants, a *Drosophila* model of neurodegeneration associated with LSD. Our results highlight an imbalance in brain ceramide and sphingosine as a crucial phenotype in the early stages of neurodegeneration. This perturbation in ceramide metabolism precedes the accumulation of endomembranous structures, manifestation of altered behavior and buildup of lipofuscin (the ageing pigment). Manipulating levels of *ceramidase*, and, consequently further altering these lipids in *spin* mutants have allowed us to conclude that ceramide/sphingosine homeostasis is the driving force in disease progression and is integral to *spin* function in the adult nervous system. Furthermore, we have identified 29 novel and direct interaction partners of Spin. We specifically focused on the lipid carrier protein, Lipophorin (Lpp), and demonstrate its localization with Spin in the adult nervous system and in organs specialized for lipid metabolism including fat bodies and oenocytes. Our observations in *spin* mutants of altered Lpp immunostaining, and of increased levels of lipid metabolites produced by oenocytes, allude to a functional relevance of the Spin-Lpp interaction.

Overall, these results detailing the kinetics of ceramide perturbations in the context of lipofuscin accumulation, as well as the proteomics experiment, represent a valuable resource to further unravel the mechanistic link between systemic changes in lipid metabolism and lysosomal storage disorders.

**Summary Statement:** Elevations in specific brain lipids and connections to relevant metabolic genes are identified in a fly model for lysosomal storage disorders. This enables a better understanding of disease progression.

## Introduction

Abnormal lipid metabolism and lipid accumulation are hallmarks of neurodevelopmental and neurodegenerative disorders. Through detailed mass spectrometry studies, perturbations in lipid metabolism have been reported [1–4]. But a systematic analysis through disease onset and progression has not yet been possible. With the help of recently established high-throughput lipidomics platforms [5,6], rapid and quantitative analyses of lipids are now feasible for complex and large studies. This has been especially useful for the characterization of lipidomes of routinely used biological model systems such as human cell lines [7], yeast [8], and *Drosophila melanogaster*. In the context of *Drosophila*, an extensively used model system, detailed lipidomics resources are now available with a focus on development and nutrition [9–11]. The fly lipidome is distinct from its mammalian counterpart in many aspects including that the major sterol is not cholesterol but ergosterol. In flies, the main membrane phospholipid is Phosophatidylethanolamine (PE) and not Phosphatidylcholine (PC) and the most abundant sphingolipid is Ceramide Phosphotidylethanolamine (CerPE), which substitutes an ethanolamine group for the choline group found in mammalian sphinogmyelins (Carvalho et al., 2012). The sphingoid-base chain length of *Drosophila* sphingolipids is shorter (predominantly C14) than in the mammalian system (C18). Furthermore, there are no polyunsaturated fatty acids (such as 20:4/ 20:5 / 22:5 / 22:6) present in *Drosophila*. Glycosphingolipids in flies constitute a core of Mannose-Glucosyl-ceramide as opposed to Galactose-Glucosyl-ceramide in mammals. Interestingly none of the reported glycosphingolipids are sialylated as in mammals, but instead have a N-Acetyl-Glucosyl-head group [12]. Despite these differences, many mammalian genes for lipid metabolism and transport are functionally conserved in *Drosophila* suggesting common principles in their metabolism and transport [13]. Furthermore, many relevant human disease associated genes connected to lipid metabolism and transport are functionally conserved; these include *Taz* mitochondrial lipid remodeling (Barth syndrome; [14], *Cerk* for ceramide metabolism (Retinitis Pigmentosa; [15] as well as *dnpc1* and *dnpc2* for lipid transport and *dsap-r*, the *pro-saposin* ortholog for sphingolipid metabolism (all lysosomal storage disorders; reviewed in [16].

Using the advancements in lipidomics, we aim to understand the nature of lipid perturbations in disease onset and progression. Contrary to other disease models or patient samples, systematic and kinetic studies on lipids are relatively easily carried out in *Drosophila,* which is a well-accepted animal model for human disease [17]. We focused on fly *spinster (spin)* mutants, which exhibit neurodegeneration with LSD-like features [16,18-20]. Lipid metabolic changes in *spin* mutants have not been characterized despite their established link to neurodegeneration and their use as genetic tool to perturb sphingolipid levels [21–23].

We have systematically analyzed and quantified lipids in adult *spin* brains at specific stages of disease onset and progression using lipidomics approaches whilst simultaneously describing the overall integrity of the nervous system using imaging, behavior, and life span analyses. Our experiments have helped establish a time line of events that place loss of *spin* function before the ceramide metabolic imbalance in the progression of disease. Using proteomics approaches coupled with pull-down assays, we have complied a resource list of Spin interactor proteins. For one of these, Lpp (Lipophorin), we present an interesting possibility that loss of Spin also causes a more general effect with changes in other lipid metabolites in tissues that regulate lipid metabolism (oenocytes).

## RESULTS

Spinster (Spin) encodes a transmembrane protein that localizes to the late endo-lysosomal compartments [20]. Early phenotypes associated with *spin* mutants include reduced viability and morphological abnormalities at the larval neuromuscular synapses and enlarged lysosomes [20]. Adult *spin* mutants exhibit signs of neurodegeneration including shortened life spans, reduced mobility, and an accumulation of electron dense material and membrane whorls in tissues [18,19]. Given the combined phenotypes of lysosomal dysfunction and neurodegeneration, these mutants constitute a good model for neurodegenerative lysosomal storage disorders. A hallmark of *spin* mutant brains is their characteristic autofluorescence that is attributed to lipofuscin or the ageing pigment. It is a mixture of accumulated/unmetabolized lipids and proteins and is distinguished by autofluorescence in ageing and degenerating tissues [24].

### Increased ceramide levels and lipofuscin in *spin* mutant brains

We imaged dissected adult brains for lipofuscin (recognized by its autofluorescent properties), and subsequently performed lipidomics on the imaged brains. In order to define the time course of lipofuscin accumulation and lipid profile alterations, we dissected brains at 1, 2, 4 and 8 days after eclosion. Our image analyses confirmed that *spin* mutant brains exhibit increased autofluorescence (Fig. 1A), which is an indication of lipofuscin [19,25]. This is in stark contrast to age matched controls that show no autofluorescence whatsoever. The heteroallelic combination *spin*^*4*^/*spin*^*5*^ was used; it displays strong *spin*-associated phenotypes for viability and synapse morphology [20] and only a limited number of escapers are viable until adulthood. These escaper adults have a reduced lifespan (Suppl Fig. 1). Lipofuscin is evident in brains as early as adult emergence (1 day) and becomes progressively more pronounced by 8 days of age. Thus, there is progressive accumulation of lipofuscin in *spin* mutants as compared to age-matched control brains. Our lipidomics analysis of single adult brains comprising of approximately 100,000 cells [26] enabled us to determine if perturbations in major membrane lipids of the brain occur in conjunction with lipofuscin accumulation. In the course of this experiment, 41 membrane lipids belonging to 6 classes were quantified. PE, PC and PE-ethers (PE-O) are the most abundant phospholipids and CerPE is the most prominent sphingolipid class in adult brains, which is similar to the larval brain lipidome [9,10,27,28].

Ceramides were the most prominently and consistently altered lipid class; an increase of over 800% on day 1 in *spin* mutant brains (Fig. 1B) as compared to controls was observed. This remarkable elevation in ceramide levels in *spin* brains was driven by the major molecular species of ceramides, 32:1 and 34:1 (Fig. 1C). It should be noted however that the magnitude of increase in ceramides in *spin* mutant brains is lowered on day 2 (over 200%), day 4 (over 350%) and day 8 (over 590%) as compared to the initial surge on day 1. The other lipid classes such as PI, PE, and CerPE show significant increases but the difference is neither sustained nor indicative of any consistent trend on the subsequent days (Fig. 1B). Because of the significant and overall increase in ceramide levels in *spin* brains at all the time-points, the remainder of our analyses has focused on sphingolipids and the early onset change on day 1

**Figure 1:**
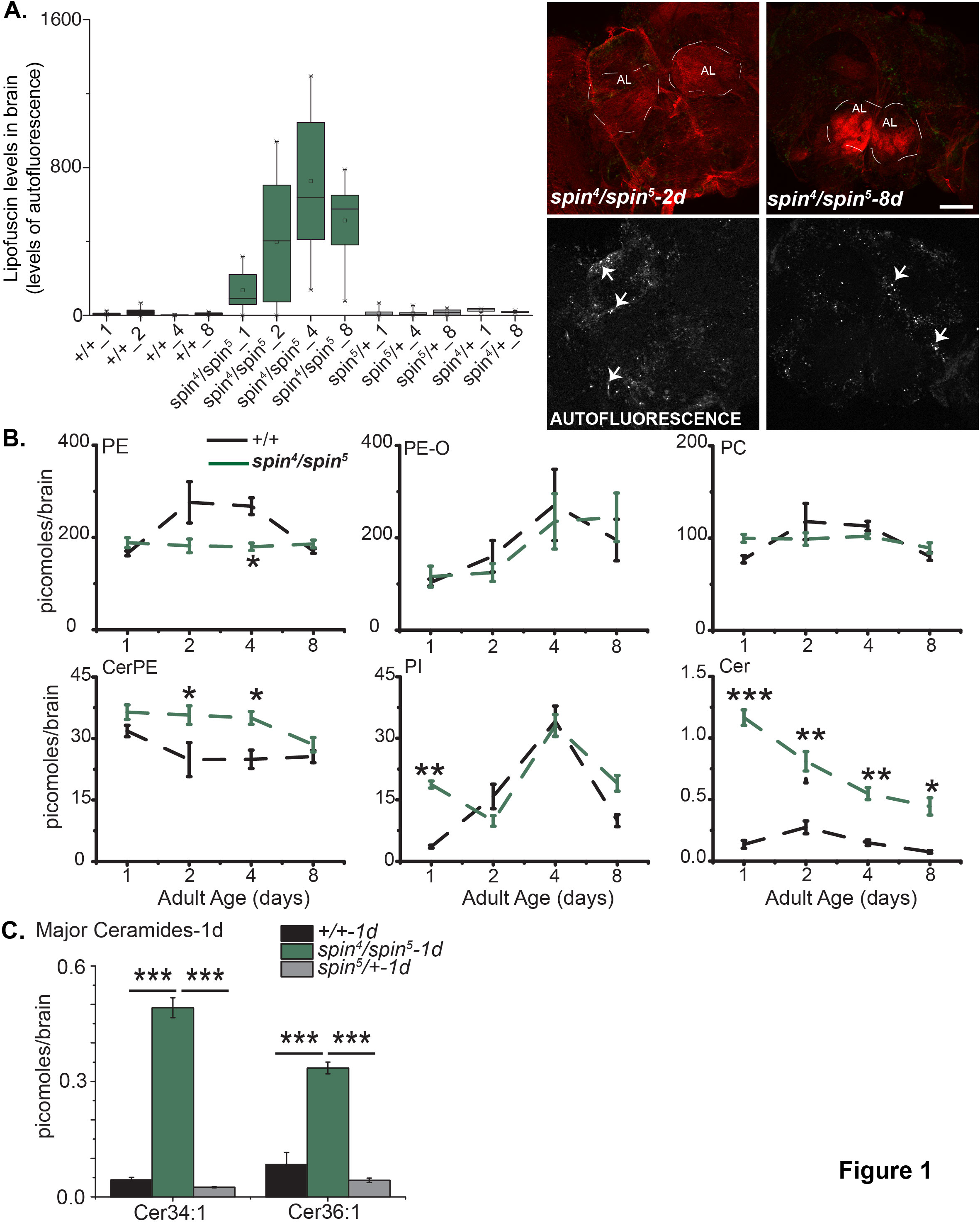
Lipofuscin accumulation and sphingolipid imbalance in *spin* mutant brains. A. Box plot represents level of lipofuscin, indicated by autofluorescence, in dissected adult brains of *spin*^*4*^/*spin*^*5*^ (green), genetic controls (*spin* heterozygotes, grey), and wild-type (+/+,black). Adult age in days is denoted after the genotype on X-axis. The box ranges between 25^th^ and 75^th^ percentile whereas the whiskers denote the minimum and maximum values in the data set. Autofluorescence levels are variable but it is evident that *spin*^*4*^/*spin*^5^ (green) brains have much higher levels of autofluorescence than age-matched genetic controls (grey and light grey) or wild-type (black) brains. Image panel (top) denotes representative images (extended field of view) of freshly dissected/unfixed *spin*^*4*^/*spin*^*5*^ brains at 2 and 8 days of age stained with Alexa 555-Phalloidin (Red). Antennal lobe neuropil (AL) are demarcated with a dotted white line. Image panel bottom constitutes the extracted green channel (greyscale) from the top images. Arrows point to autofluorescent puncate and are indicative of lipofuscin. Scale bar = 100mm B. Values are mean ± S.E.M. for lipid levels (quantified as picomoles/brain) from single brains previously imaged for lipofuscin (Top panel). Dotted lines are indicative of trends over the ages analyzed (1,2,4 and 8 days of age). Green indicates *spin4/spin5* whereas black indicates wild-type (+/+).*p<0.005, **p<0.00005, ***p<0.00000005 as indicated by post-hoc Tukey test following ANOVA C. Bar graphs represent absolute levels of major ceramide species (Cer34:1 and Cer36:1) on Day 1 in brain extracts of *spin*^*4*^/*spin*^*5*^ (green), genetic controls (*spin* heterozygotes, grey), and wild-type (+/+,black). Values are mean ± S.E.M. quantified as picomoles/brain. ***p<0.00000005 as indicated by post-hocTukey test following ANOVA

### Occurrence of lipofuscin correlates with phenotypic severity of *spin* mutants

The occurrences of lamellated whorls and endomembranous structures are characteristic ultrastructural hallmarks of several degenerative diseases including LSD. Indeed such structures have been previously reported in *spin* mutant retinas and in other fly models of LSD [18,19,29]. We therefore asked if ultrastructural abnormalities in *spin* mutant brains coincide with altered lipid levels. Three categories of abnormal structures were observed in *spin* mutant brains that were not observed in age-matched control brains (Fig. 2A); these include lamellated whorls/multi-lamellar whorls, prominent electron dense lysosomal structures, and a combination of dense bodies with lamellated whorls and membrane bound structures (Fig 2B, left to right). The last category is reminiscent of lipofuscin - an outcome of increased oxidative stress and intracellular debris in degenerating and ageing cells. Lamellated whorls and prominent lysosomes were observed at a higher frequency at the onset (1-2 days of adulthood) as compared to 8 days, when lipofuscin-like structures become increasingly abundant (Fig. 2C). These structures are also consistently observed in another allelic combination, *spin*^4^/*spin*^Δ*58*^, that exhibit a milder phenotype with regards to adult life span (Suppl Fig 1). When we compared the two allelic combinations, the more severe *spin*^*4*^/*spin*^*5*^ versus the milder *spin*^4^/*spin*^Δ*58*^, we recognized that the stronger phenotype with reduced life span (Suppl Fig 1) correlated with an earlier occurrence and a higher frequency of lipofuscin (Fig. 2C). It should be noted that these abnormal structures were not observed in age-matched control brains at these early stages of the adult lifespan i.e. up to 8 days after eclosion. We conclude that because lamellar whorls precede the formation of lipofuscin, they constitute an earlier phenotype than lipofuscin in the disease progression.

**Figure 2:**
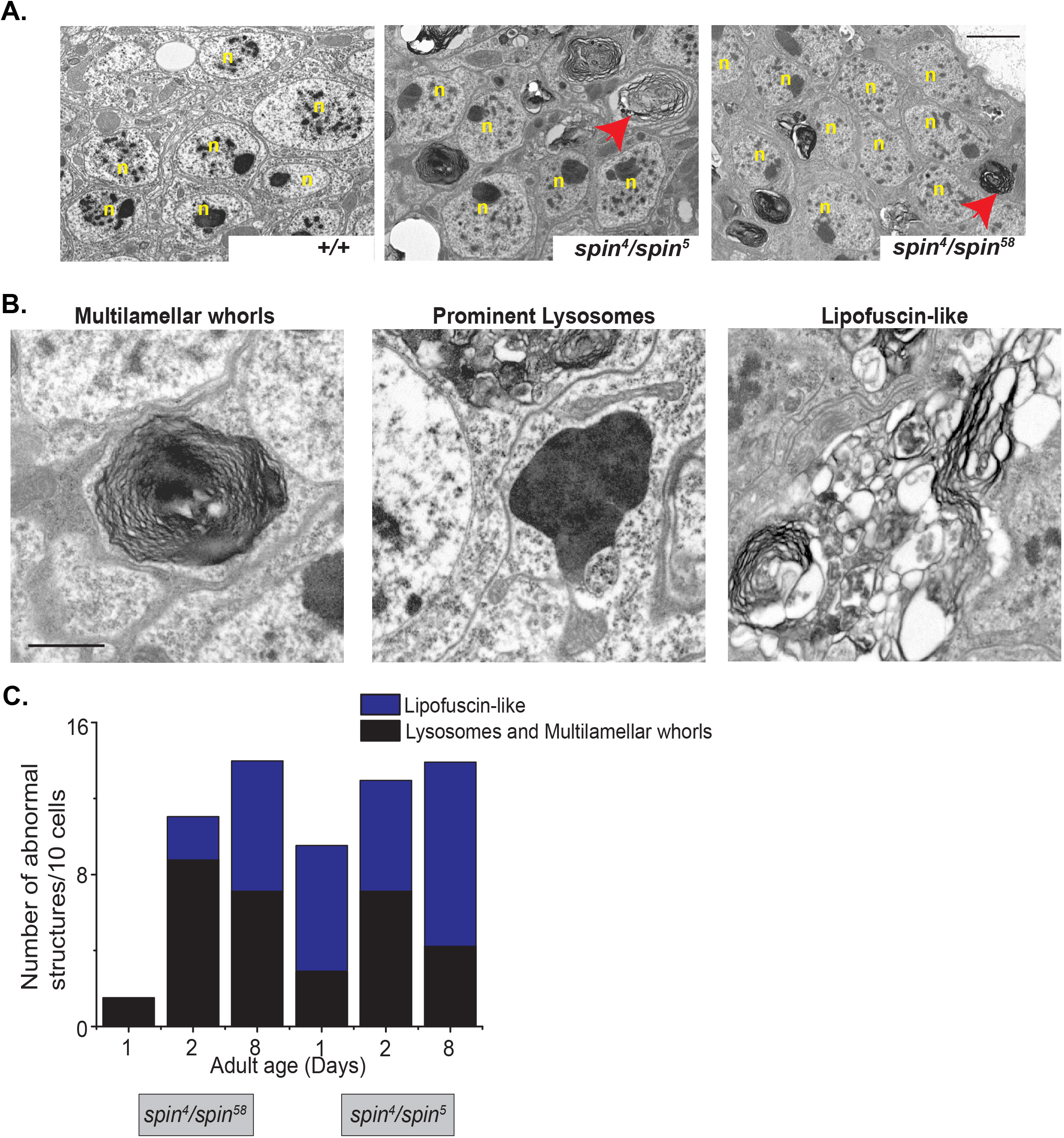
Abnormal accumulation of endomembranous structures. A: Representative TEM images of cell bodies in the cortex of the adult brain at 2days for *spin*^*4*^/*spin*^*5*^, *spin*^*4*^/*spin*^58^, and wild-type (+/+). In both *spin* mutant heteroallelic combinations abnormal structures (red arrows) were observed. Nuclei of the cell bodies are indicated as n, Scale bar =2µm B: Representative images of different endomembranous structures observed in *spin* mutant brains; (i) lamellar whorls, (ii) electron dense lysosomes and (iii) lipofuscin. Scale bar =1µm C: Occurrence of abnormal structures (lipofuscin, lamellar whorls and prominent lysosomes) quantified as number of abnormal structures normalized to 10 cells (n=3 animals) in *spin* mutant transheterozygotic (*spin*^*4*^/*spin*^*5*^, *spin*^*4*^/*spin*^*58*^) brains. In age-matched wild-type brain (1-8 days of adulthood), these abnormal structures are almost never observed.

### Degree of ceramide and sphingosine elevation in adult *spin* brains correlates with the severity of neurodegeneration

The increase in ceramide levels, observed with single brains (Fig. 1), was further confirmed when we measured ceramide and its low-abundant breakdown metabolite, sphingosine, using larger amounts of brain tissue. Consistent with single brain measurements, we observed a dramatic increase in ceramides on day one of adulthood (Fig. 3A) in *spin*^*4*^/*spin*^*5*^ mutants. For sphingosine, we observed a tremendous increase in abundance for *spin* mutants (Fig. 3). We also note a smaller but significant increases in sphingosine and ceramide levels in the second heteroallelic combination *spin*^*4*^/*spin*^*Δ58*^ but to a lower extent (Fig. 3). Here, for ceramides, statistically significant changes were only detected in ceramide dienes. Overall, the changes for *spin*^*4*^/*spin*^*Δ58*^ are less pronounced when compared with *spin*^*4*^/*spin*^*5*^ brains but are consistent with the milder survival and ultrastructural phenotypes.

**Figure 3:**
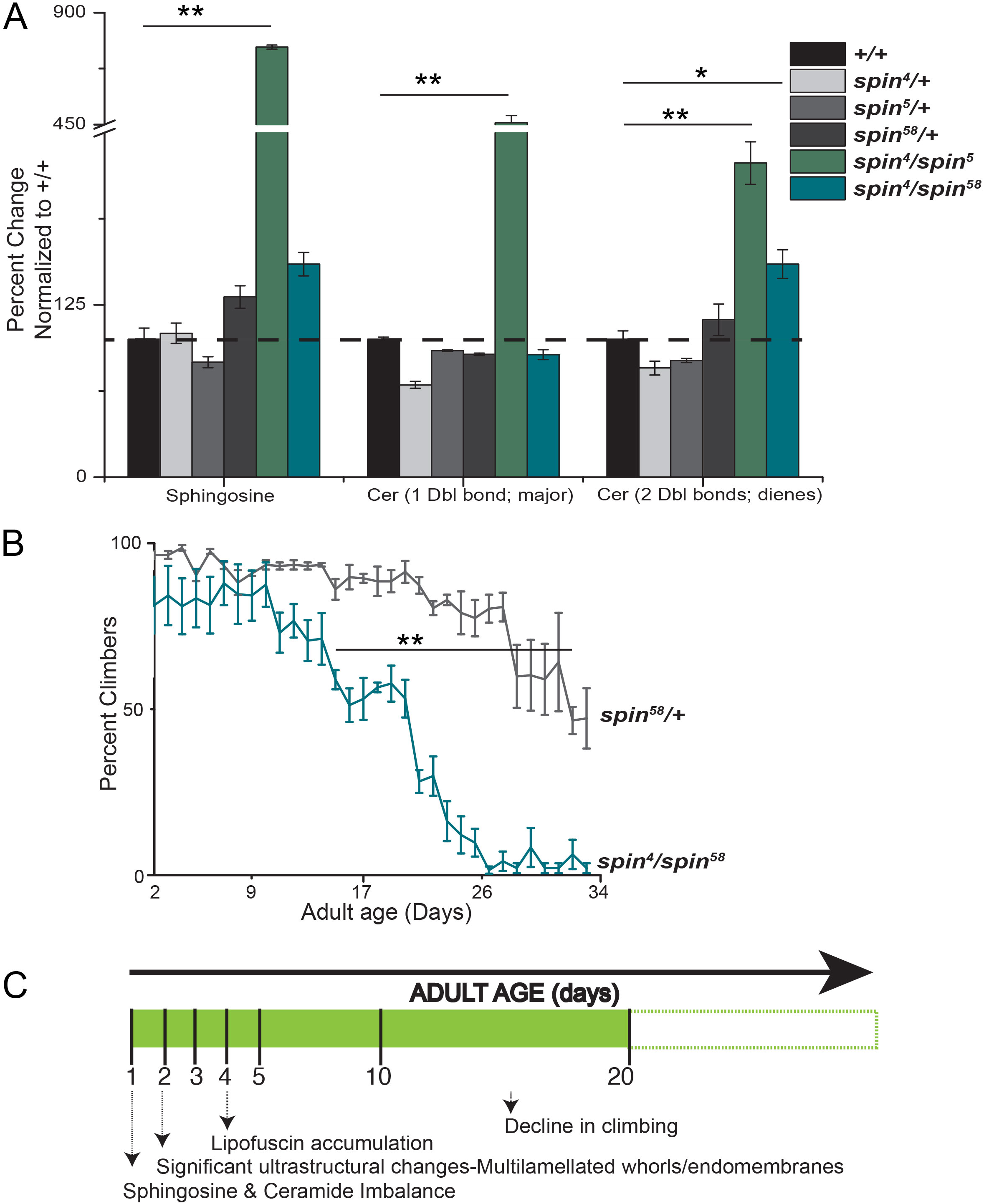
Ceramide/Sphingosine imbalance occurs before behavior change. A. Altered sphingosine/ceramide levels in two heteroallelic combinations of *spin* mutants. Bar graphs indicate percent change in levels of sphingosine (Sph) and ceramides (1 double bond or 2 double bonds; diene species) measured on day 1 of eclosion in adult brains compared to wild type. Values are mean± S.E.M., and *p<0.05 and **p <0.01 as calculated by post-hoc test following ANOVA. Black line indicates the baseline levels in control (+/+) to which all other lipid levels (picomoles/brain) are normalized. B. Progressive loss of climbing behavior in *spin* mutant. Line graphs indicate trend of changing climbing behavior; each data point constitutes the daily average for percent climbers ± S.E.M. from three independent trials of 10-15 flies/trial over age (X-axis). **0.001<p<0.05, Student’s t-test. C. Timeline relating ceramide and sphingosine imbalance during adult lifespan and the onset of specific neurodegenerative attributes such as lipofuscin accumulation and decline in climbing behavior in *spin* mutants.

The enhanced viability of *spin*^*4*^/*spin*^*Δ58*^ heteroallelic combination over *spin*^*4*^/*spin*^*5*^ enabled us to investigate the chronological coincidence of ultrastructural abnormalities, lipid alterations and behavioral impairment. Here, we used changes in the climbing ability of adult flies as a general indicator of behavioral alteration/impairment. Decreased climbing ability is displayed only after two weeks of adult age (Fig. 3B; see Movies in supplemental section). Our experiments suggest that ceramide and sphingosine metabolic perturbations constitute an early-onset event in these mutants, after which other signs of neurodegeneration such as ultrastructural abnormalities, lipofuscin accumulation, and behavioral impairment become pronounced (time line summarized in Fig. 3C).

### Ceramide imbalance is central to the role of *spin* in maintenance of adult neurons

To show that *spin* function is causally connected to perturbed ceramide/sphingosine levels, we carried out a genetic rescue experiment wherein full-length wild-type *spin* is overexpressed in the background of the mutant, *spin*^*4*^/*spin*^*5*^, specifically in cells where the endogenous *spin*-promoter is active [20]. Rescued animals have a life span similar to controls with a 50% survivorship (T-50) of 45 days as compared to 30 days in the mutants (Suppl Fig 1). Lipid metabolic alterations are also rescued by the transgene in this background. The abnormally high levels of brain ceramide and sphingosine in *spin*^*4*^/*spin*^*5*^ were reduced in rescued genotypes to levels comparable to the controls (Fig. 4A). Finally, the ultrastructural hallmark of *spin* mutant brains, which is the abundance of lamellar whorls, is significantly diminished upon genetic rescue with *spin* overexpression (Fig. 4B). Thus sphingolipid imbalance and the associated neurodegenerative attributes (reduced survival, progressively shortened lifespan, and ultrastructural abnormalities) described for *spin* mutants are all rescued by an overexpression of *spin*.

**Figure 4:**
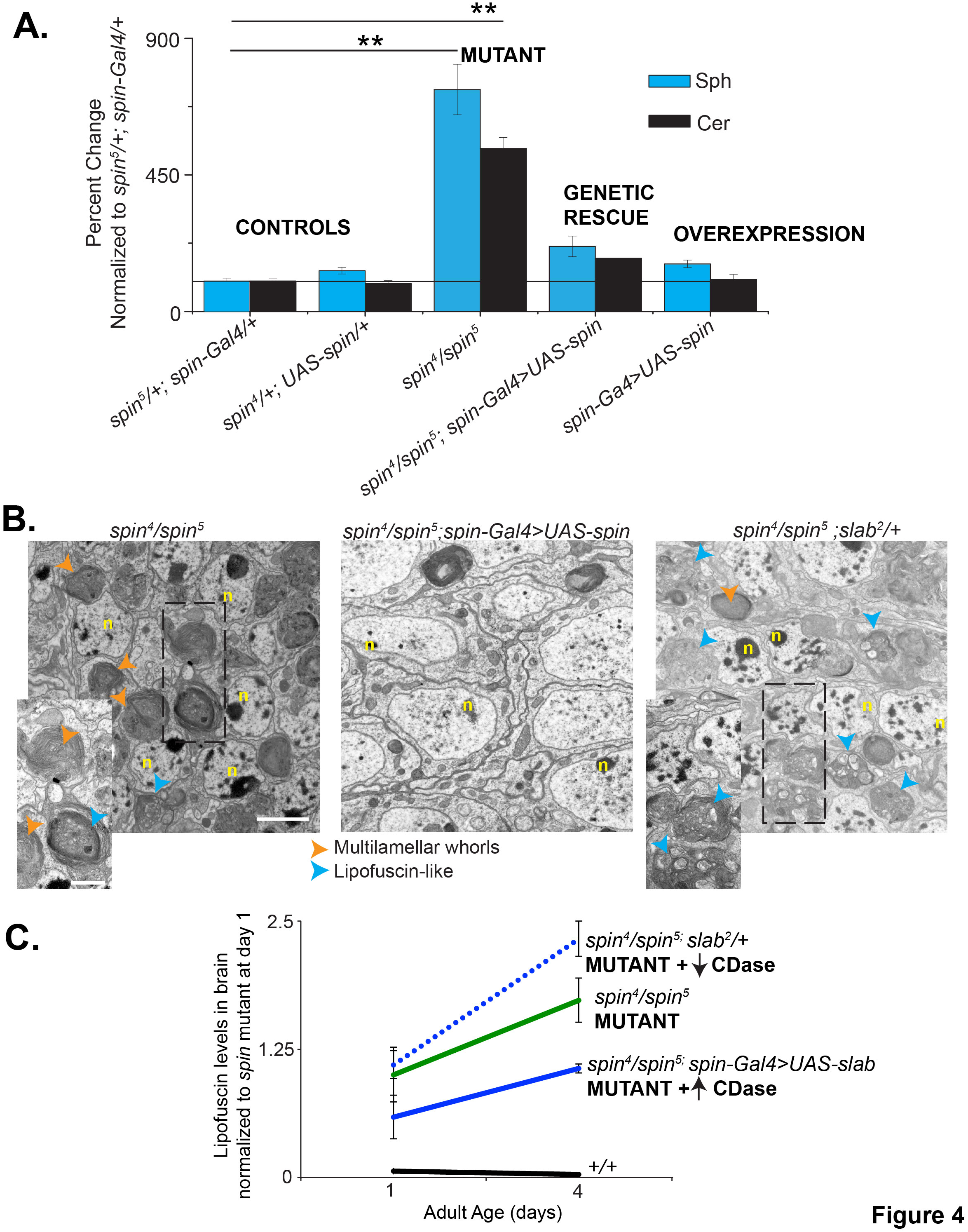
Rescue of *spin*^*4*^/*spin*^*5*^ phenotypes with *spin* overexpression and modification of *spin*^*4*^/*spin*^*5*^ phenotypes with *slab (Ceramidase)* manipulation. A. Bar graphs indicate percent change in levels of sphingosine (blue bars) and total ceramides (black bars) measured on day 1 of eclosion in adult brains for mutant, genetic controls, rescue and overexpression genotypes (also indicated along the X-axis). Lipid levels are quantified as picomoles of lipid/brain and percent change with respect to genetic control (*spin*^*5*^/*+; spinGal4/+*) are calculated, values are mean± S.E.M., and **p<0.000001 as calculated by post-hoc test following ANOVA. B. Representative TEM images of cell bodies in the cortex of the adult brain at 4 days of *spin*^*4*^/*spin*^*5*^ (mutant), *spin*^*4*^/*spin5; spin-Gal4>UAS-spin* (genetic rescue) and *spin*^*4*^/*spin*^*5*^; *slab*^*2*^/*+* (mutant + decreased *Cdase*). In *spin* mutants alone and mutant + decreased *Cdase* combinations, abnormal structures (arrows) were observed. Insets show these abnormal structures at a higher magnification. In case of the mutant alone, there are prominent lamellar whorls (orange) and a few lipofuscin-like strunctures (blue). These abnormal structures are strikingly absent upon genetic rescue but in case of mutant + decreased *Cdase* combinations, there is an abundance of lipofuscin-like strunctures (blue). Nuclei of the cell bodies are indicated as n, Scale bar =1.8µm and 0.8µm for insets. C. Line profiles represent the mean ± S.E.M. for normalized level of lipofuscin (indicated by autofluorescence) at 1 and 4 days of age in dissected adult brains of *spin*^*4*^/*spin*^5^ (green), wild-type (+/+,black), and upon manipulation of *slab* (Cdase) in the background of *spin* mutants-*spin*^*4*^/*spin*^*5*^; *slab*^*2*^/+(blue dotted) and *spin*^*4*^/*spin*^5^; *spin-Gal4> UAS slab* (blue). The levels are normalized to that of *spin* mutants on day 1 of eclosion. Lipofuscin levels steadily increase in *spin* mutants between day 1 and day 4. Introduction of *slab/+* in the *spin* background results in an even greater accumulation of lipofuscin whereas this trend is reversed with an overexpression of *slab* in the *spin* mutant background.

To test if ceramide/sphingosine imbalance is causal to the degenerative attributes in *spin* brains, the *slab* (*ceramidase /Cdase;* [30] gene was manipulated in the background of *spin* mutants. Mutants and overexpression lines of *slab* exhibit altered levels of ceramides as characterized by MS measurements [31,32]. *slab* heterozygotes and overexpression of *Cdase* specifically in neurons showed an increase and decrease of brain ceramides by over 200% and 30% respectively (Hebbar et al, 2015). Therefore, these genetic tools were combined in the background of *spin* mutants. Ultrastructural defects as well as lipofuscin accumulation were used to score the extent of the degenerative phenotype against *spin* mutant alone. Reducing *slab* levels (with *slab*^*2*^ heterozygosity; *slab*^*2*^/*+*) in the background of *spin* mutant, and consequently further pushing ceramide levels higher (documented for *slab*^*2*^/*+* in [33], correlated with an intensification of these degenerative phenotypes. This effect is most clearly evidenced (at day 4 of adulthood) by the increased presence of ultrastructural defects (Fig. 4B) and enhanced autofluorescence by 34% (Fig. 4C) in *spin*^*4*^/*spin*^*5*^; *slab*^*2*^/*+*. The abundance of lipofuscin-like structures as opposed to lamellar whorls in age-matched *spin*^*4*^/*spin*^*5*^; *slab*^*2*^/*+* neurons (Fig. 4B) highlights the severity and progression of degeneration caused by a reduction of *slab* function (and increase in ceramides) in *spin*^*4*^/*spin*^*5*^ brains. In contrast, overexpression of *slab*, and consequently reducing ceramide levels (documented in [33], halted the formation and accumulation of lipofuscin with age (Fig. 4C). A 42% reduction in lipofuscin levels was seen at the onset (day 1) and a 39% reduction was observed at a later stage (day 4) in *spin*^*4*^/*spin*^*5*^; *spin-Gal4>UAS slab* as compared to *spin*^*4*^/*spin*^*5*^. This epistatic relationship between ceramide manipulation and *spin* mutant abnormalities suggests that lowered ceramide levels in *spin* brains correspond to lowered accumulation of lipofuscin. However it should be noted that *slab* manipulation did not correlate with developmental lethality/viability of *spin*^*4*^/*spin*^*5*^; *slab* overexpression and *slab*^*2*^/*+* in *spin*^*4*^/*spin*^*5*^ both caused increases in *spin*^*4*^/*spin*^*5*^ viability (of 7% and 20% respectively) to adulthood. This finding probably indicates that the genetic interaction of *slab* and *spin* is not relevant for the role of *spin* in overall viability.

The *spin* rescue experiments and the genetic interaction with *slab* help delineate that the removal of *spin* function in *Drosophila* causes an early-onset imbalance in sphingolipid metabolism during adulthood. Elevated levels of ceramides and sphingosine are responsible for the progression of the disease (specifically of lipofuscin accumulation) leading to a loss of neuronal health and to changes in behavior. It remains unclear if these lipids are specifically enriched in the observed ultrastructural abnormalities.

### Interactors for Spin have roles in carbohydrate and lipid metabolism

In an effort to identify intracellular processes impaired by loss of function of *spin*, we sought to determine its direct interactors by co-immunoprecipitation and proteomics (see Suppl Methods-Schematic 1 for details). In this way, 29 interactors for Spin were detected and are summarized in Table1. Using the PANTHER Classification System [34,35], these proteins were classified according to their listed cellular component ontology. Thirteen proteins localize to the mitochondrion and are involved in glycolysis. Glycolytic enzymes are recruited to the synapses during energy demand [36]. Since Spin is actively trafficked along axons [37,38] via an energy demanding process, these proteins constitute a potentially intriguing cluster. Further, one protein (Guanine nucleotide-binding protein subunit beta-like protein/Rack1) is assigned to the endosome whereas the lipoprotein (Apolipophorin; ApoLpp) is assigned to the autophagosome; both these sub-cellular locations are relevant given the previously published information on Spin [20,38,39]. In the context of a multi-organ and systemic lipid metabolic imbalance in LSD, we focused on the relevance of Lipophorin (Lpp; the functional lipoprotein produced from ApoLpp) as a direct interactor of Spin. Lpp constitutes the lipid transport protein in *Drosophila* that originates in the fat bodies and facilitates inter-organ lipid transport including the delivery of lipids [28,40]. In order to confirm the interaction between Lpp and Spin, we investigated Lpp and Spin localization in tissues throughout the body. In the adult brain, Spin-GFP driven by *spin-Gal4* is seen in the neuropil and cortex. In the cortical areas, punctate staining for Spin is evident and a subset of these structures are positive for Lpp immunoreactivity (Fig. 5A-A”). We further observed co-localization in fat bodies, where both Spin-GFP and Lpp [40] are also abundantly present (Fig. 5B). Consistent with neuronal staining, a subset of Spin and Lpp are co-localized in compartments that are in close proximity to the membrane (Fig. 5B’).

**Figure 5:**
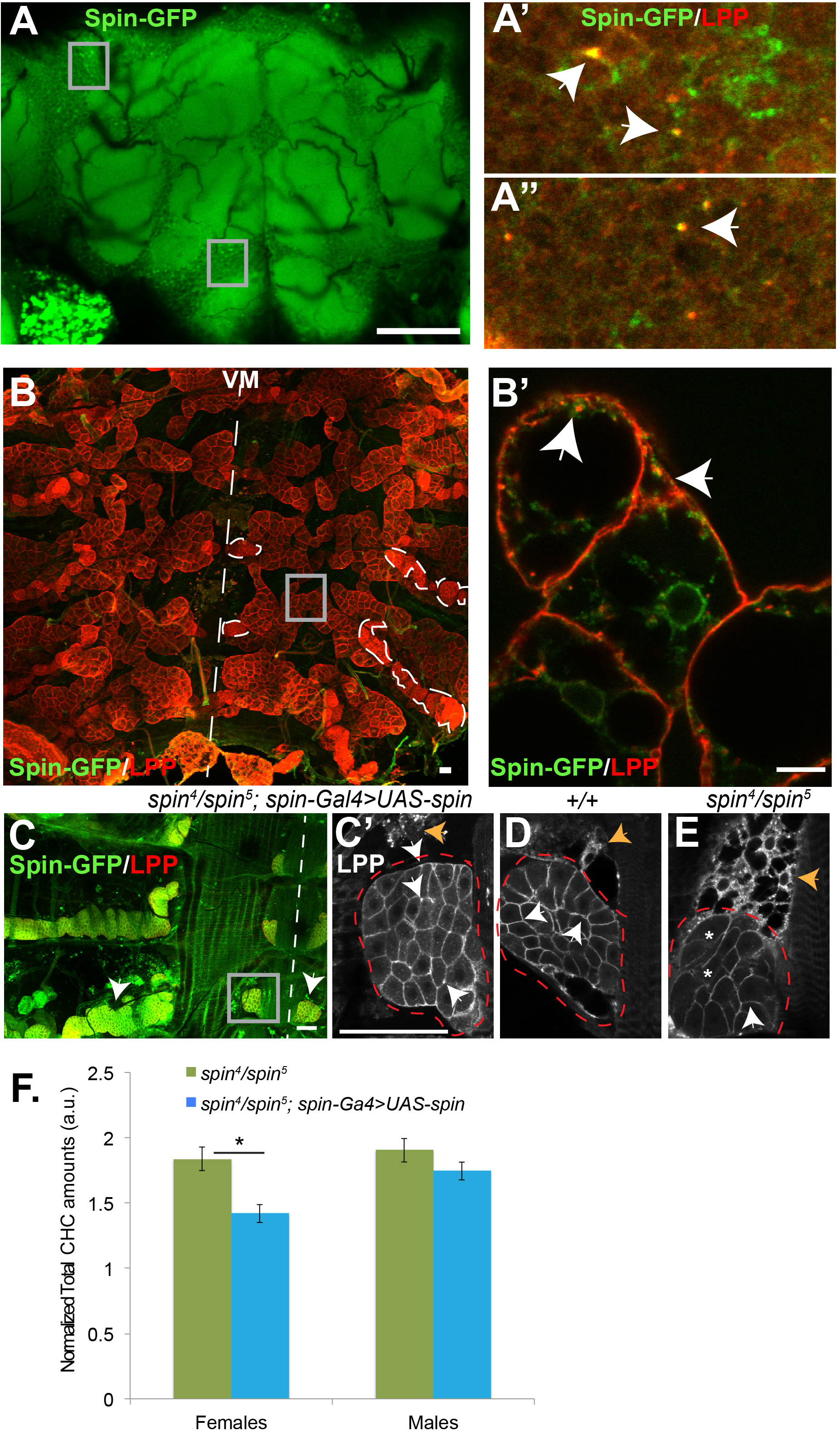
Spin-Lpp Interaction. A-B’: Representative images of (A-A”) adult brain and (B-B’) adult abdomen of *spin-Gal4>Spin-GFP* animals labeled with GFP (Green) and Lpp (red) antibodies. Scale bar =50µm. In the abdominal fillet preparation, ventral mid-line is indicated with a dashed white line and fat bodies (grey box) and oenocytes (dashed outline) display LPP staining. Boxed regions are imaged at high magnification and shown in A’-A”, B’. Arrowheads indicate subset of Spin-GFP-positive punctae that also display LPP immunoreactivity in neuronal cortex (above) and in fat bodies (below). Scale bar: 10 µm. C: Female adult abdominal fillet labeled with GFP and Lpp antibodies in rescued (*spin*^*4*^/*spin*^*5*^; *spin-Gal4>UAS-spin)* genotype. Rescue construct (Spin-GFP; green-C) is clearly evident in dorsal and ventral oenocytes (arrows in C); Ventral mid-line is indicated by white dashed line. Scale bar: 50 µmLpp distribution in ventral oenocytes (red outline) and adjoining fat bodies (orange arrow) is shown at a higher magnification (C’-E) in rescue (C’; *spin*^*4*^/*spin5; spin-Gal4>UAS-spin*), wild-type (D; *+/+*) and in mutant (E; *spin*^*4*^/*spin*^*5*^) genotypes. Lpp is visible at the membrane of oenocytic cells (white arrows) but the intensity of Lpp staining is reduced in *spin* mutants (E); there are some clusters of cells (white asterisk) wherein the membrane staining of Lpp apprears diminished. Scale bar: 50 µm F. Bars represent mean ± S.E.M. for total cuticular hydrocarbon (CHC) levels in male and female flies from mutant (*spin*^*4*^/*spin*^5^) and (*spin*^*4*^/*spin5; spin-Gal4>UAS-spin*) rescued genotypes. Mutant females display significantly (p<0.05) higher levels of CHCs as compared to rescued genotypes; p=0.01, post-hoc Holm-Sidak test following ANOVA; n=3 replicates of 6-8 flies each.

Interestingly in *spin* rescue animals, oenocytes are labeled very strongly when Spin-GFP is driven using *spin-Gal4* (Fig. 5C). Adult oenocytes are a band of cells present dorsally and ventrally along the abdominal cuticle (Fig. 5B, C and Suppl. Fig. 3; [41]. Oenocytes regulate lipid metabolism and energy storage [42]; making them an ideal tissue to gain further functional insight into the Spin-Lpp connection. Adults oenocytes act as centers for the production of cuticular hydrocarbons, several of which function as pheromones [41,43]. Since cuticular hydrocarbons (CHCs) are synthesized by the extension and modification of fatty acid precursors [43], we hypothesize that Spin-LPP will influence CHC synthesis in the oenocytes. Support for this hypothesis comes from two observations. First, Lpp immunostaining in *spin* mutant oenocytes (Fig. 5E) is altered, with a reduced intensity (as compared to rescue conditions (Fig. 5C’) and wild-type conditions (Fig. 5D); and more examples in Suppl Fig.3). Second, total amounts of CHCs are significantly increased in *spin* mutant females (Fig. 5D) whilst there is no change in specific CHC content. Taken together these observations point to an intriguing possibility that Spin-Lpp interaction plays a role in regulating CHC synthesis. Whether it is the delivery of the fatty acid precursors to the oenocytes, or the actual biosynthesis, or the subsequent transport of the CHC from the oenocytes, remains to be addressed. Moreover whether more general deficits in lipid metabolism and inter-organ lipid transport occur, is an interesting question for future research.

## Discussion

We identified alterations in brain ceramide and sphingosine levels in adults following loss of *spin* function. Perturbations in lipid quantities occur prior to the onset of endomembrane accumulation, and before the functional and morphological attributes of neurodegeneration are evident (summarized in Fig. 3C). These observations are comparable with reported elevation in sphingosine in cell lines upon drug-induction of LSD [44]. A recent study has linked oxidative stress, lipid droplet formation, and neurodegeneration in *Drosophila* and mice tissues [45]. Our approach has enabled us to define the nature and dynamics of these lipid perturbations and confirm the suspected links of lipid imbalance and pathological attributes of degeneration in a temporal and tissue-specific manner.

We argue that this loss of ceramide/sphingosine homeostasis is not due to apoptosis but is directly linked to loss of *spin* function. An indication that enhanced apoptosis is not the underlying cause is evident from the absence of any notable apoptotic-like features (vacuolization) in the TEM images of 1-day-old brains (Fig 2). That *spin* function is causally connected to ceramide and sphingosine is further validated by our rescue experiments.

The *slab*/*Cdase* and *spin* interaction experiments clearly establish loss of *spin* function upstream of the ceramide imbalance. This alteration in sphingolipid levels is responsible for lipofuscin accumulation. Given the chronology of events (summarized in Fig. 3C) and the outcome of our genetic interaction/rescue experiments, it is clearly evident that the accelerated lipofuscin accumulation, that is a result of perturbed ceramide levels, correlates with the loss of nervous system integrity in *spin* mutants. How does the loss of Spin result in a perturbation of sphingolipids? One possibility is linked to the previously reported function of Spin as a Sphingosine-1-Phosphate transporter [21]. Since our measurements are limited snapshot measurements of brain lipids, we cannot make any speculations on the dynamics of ceramide/sphingosine and S1P transport in relation to *spin* function. It is possible that the loss of Spin causes an altered partitioning of S1P in the late endosomal/lysosomal compartments, which leads to aberrant signaling. A second likely possibility stems from observations that loss of Spin results in the incomplete progression of lysosomal reformation and the accumulation of autolysosomes [39]. We hypothesize that altered *spin* function in adult neurons accelerates or enhances the formation of autophagic structures such as autolysosomes. This disruption leads to a failure to completely recycle components including lipids such as ceramides and sphingosine. Since the process is most likely stalled, and further exacerbated by the elevated levels of these signaling lipids, there is an eventual accumulation of intracellular debris in the form of lipofuscin. In motorneuron degeneration of *bchs* mutants, perturbations in autophagic flux cause the accumulation of these signaling lipids in autophagosomes, and this prevents pro-survival signaling involving MAP and Akt [33]. Here, our EM data effectively demonstrates that changing levels of ceramide has consequences on disease progression by causing a detrimental accumulation of lipofuscin in *spin* mutants. It is therefore very likely that in a similar fashion to *bchs* larval motorneuron degeneration an overlap between perturbations in ceramide/sphingosine signaling and *spin* function in autophagy underlie the age-associated degeneration of adult *spin* mutants. Since all these three processes namely a lipid metabolic imbalance, neurodegeneration and autophagy are implicated in LSD, this work extends our understanding of the sequence of events in this disease model. Some relevant aspects of the above model are however unresolved. First, it will be interesting to elucidate the nature of ceramide/sphingosine downstream signaling that causes the aberrant progression of autophagy and leads to the formation of lipofuscin. Likewise, it will be important to also address the exact roles played by sphingosine and ceramide and if an imbalance is only limited to signaling in/from autophagic structures. Indeed our experiments with *slab* and *spin* suggest that changing overall levels, presumably in all membranes, certainly has an impact. Therefore it will be important in the future to use approaches that enable manipulation of these lipids in specific sub-cellular compartments and monitor the consequences on disease progression.

It is known that *spin* directly or indirectly regulates important signaling cascades including TGF-ß/bone morphogenetic protein [20] and JNK/AP-1 (Milton et al., 2011) at the synapse. Further a close association between lipid metabolic gene disruption and altered morphogen signaling [23] has been previously reported. It remains to be seen if an overlap between *spin* function, lipid metabolism, and morphogen signaling has implications in synapse expansion.

We have also identified potential metabolic and cellular interaction partners of Spin including an interaction with Lpp. Our observations point to the likelihood that Spin and Lpp interact in adult oenocytic function, specifically in the overall production/transport of CHCs. It also raises the hypothesis that Spin/Lpp interaction may be relevant in inter-organ lipid transport routes and could have implications in systemic diseases such as LSD where multiple organs are affected. In conclusion, studying the contribution of these interactors will throw light on the increasingly acknowledged connections between lipid metabolic imbalance, loss of cellular integrity, and the manifestation of disease pathology.

## Materials and Methods

### Flies

All fly lines and genotypes used are listed in Supplementary Methods; Suppl Table 1. Flies were raised at 25°C on a regular yeast-sugar-cornmeal medium supplemented with fungicidal mixture of phosphoric acid and propionic acid (http://flystocks.bio.indiana.edu/Fly_Work/media-recipes/caltechfood.htm).

### Lipofuscin Imaging and Analyses

Adult brains were dissected in phosphate buffered solution on ice. They were placed in Alexa568-labelled Phalloidin (1:40; Life Technologies, India) for 20 minutes on ice before transferring onto PBS on glass slides for imaging. Brains were imaged using Fluoview 1000 (Olympus Corporation, Japan) with an Argon 488 (autofluorescence) and HeNe (568 Phalloidin) laser. Optical sections were taken and merged using Image J [46] to create an image stack. Image stack was analyzed to give particle count to indicate number of fluorescent particles.

### Immunostaining

Dissected adult brains and abdominal fillets were immunostained following [47]. The primary antibodies used were anti-GFP (A-11122, 1:200; Life Technologies, India) and Anti-Lpp (guinea pig, gift from Marko Brankatschk; [40]. Fluor-tagged secondaries were obtained from Life Technologies, India.

### Transmission Electron Microscopy

Adult heads were dissected, placed in fixative (4% paraformaldehyde, 1% glutaraldehyde in 0.1M sodium phosphate buffer, pH 7.4) and vacuum treated to removed adherent air. Heads were incubated in fresh fixative overnight at 4°C. All incubations were on a rotating wheel, unless otherwise stated. Heads were washed in 0.1M sodium phosphate buffer (3 x 10 min), post-fixed in 1% OsO_4_ for 1 h, followed by washes in 0.1M sodium phosphate buffer (3 x 10 min) and dH_2_O (3 x 10 min). Heads were dehydrated in an acetone series (30%, 50%, 70%, 90%, 3x 100%; 20 min each) then incubated in increasing concentrations of Spurr’s resin:acetone (25%, 50%, 75%, 95%, 2x 100% [at 37°C]; 45 min each) followed by incubation in 100% resin overnight at 4°C without rotation. Heads were embedded in Spurr’s resin for 24 h at 70°C.

After reaching the desired depth by semi-thick sectioning, ultrathin sections (60–70 nm) were collected coated grids, treated with uranyl acetate in 50% ethanol (10 min) and submerged in dH_2_O to wash. Sections were stained with lead citrate (10 min) in the presence of sodium hydroxide pellets, followed by washing in dH_2_O. Images were acquired using analysis software on a TECNAI G^2^ (Version 2.18) transmission electron microscope (120 kV).

### Lipid Extraction and Lipidomics

#### Lipid extraction

Female adults of the required genotypes were collected on emergence and aged appropriately. Depending on the experiment, single brains or groups of 3 brains were dissected in PBS and flash frozen in liquid nitrogen and stored in 20% methanol to avoid any enzymatic degradation. Brain samples were homogenized and extracted according to the methyl-tert-butyl ether extraction method [48]. All lipid standards were added to the homogenates prior extraction (Suppl Table 2). The upper MTBE layer was dried and stored until measurement at −80C. For the mass spectrometric measurements, the samples were dissolved in 100µl methanol containing 0.1% ammonium acetate.

#### Lipidomics of single brains

To study the correlation between lipfuscin accumulation and lipid metabolism (Fig. 1), single brains were homogenized directly after imaging and extracted with the MTBE based extraction protocol. LC-MS was performed on a Agilent 1200 micro-LC system (Agilent Technologies, USA) coupled with a LTQ Orbitrap XL mass spectrometer (Thermo Fisher Scientific, Bremen, Germany) using a split flow setup to perform nano-ESI with the Nanomate TriVersa (Advion BioSciences Ltd, USA). Liquid chromatography was performed at a flow rate of 20µl/min using a Zorbax SB-C18 column (0.5mm ID, 5µm, 150mm) and 5µl sample were injected. Solvent A was methanol containing 0.1% ammonium acetate and solvent B MTBE. The gradient was as follows: 0-3 min, B = 0.5%, 18–21 min B = 50% and 22–37 min B = 0.5%. Extracted ion chromatograms for designated lipids were integrated using accurate masses (± 5ppm) with Xcalibur software (Thermo Fisher Scientific, Germany)

#### MS^n^ analysis of pooled brain lipid extracts

Lipid extracts were analyzed with a flow-injection system using a flowrate of 1µl/min and 5µl sample injection. Negative and positive ion mode spectra were acquired with a LTQ-Orbitrap XL equipped with a 1200 micro-LC system and a Nanomate Triversa utilizing 5µm ID ESI-chips. In the negative mode PI, PE, PE-O, LPE, PC, CerPE, PS, PG were identified according to their accurate mass as described earlier [5]. For PS, the specific neutral loss of 87Da was monitored in the linear ion trap and used for quantification. In the positive ion mode Sph 14:1, Ceramides and HexCeramides were monitored with MS^3^ in the linear ion trap using the long chain base related fragment ions. All MS^3^ for quantifying sphingolipids were analyzed using Xcalibur software while all other analyses were performed using LipidXplorer [49].

For single and pooled brains, absolute levels of individual lipid species were summed up to arrive at lipid class quantities. At least 5 biological replicates were used for the analyses. GraphPad Prism (GraphPad Software, Inc, USA) and Origin 8.1 (OriginLab, USA) were used to for graphical representation and Origin 8.1 for ANOVA analysis coupled with post-hoc Tukey test.

#### Annotation of Lipids

Phospholipids were indicated as ⟨lipidclass⟩ ⟨no. of carbons in all fatty acids⟩:⟨no. of double bonds in all fatty acids⟩. For the annotation of phosphatidyl ethanolamine vinyl ether and alkyl ether the abbreviation (PE-O) was used along with the number of double bonds of the fatty acid and fatty alcohol moiety. Sphingolipids were annotated as ⟨lipid class⟩ ⟨no. of carbons in the long-chain base and fatty acid moieties⟩:⟨no. of double bonds in the long-chain base and fatty acid moieties). Lipid class abbreviations were utilized throughout the manuscript as follows: phosphatidyl ethanolamine (PE), phosphatidylethanolamine ether (PE-O), phosphatidylcholine (PC), phosphatidylinostiol (PI), ceramide (Cer), phosphatidylserine (PS), ceramide phosphorylethanolamine (CerPE), Sphingosine (Sph).

#### Co-Immunoprecipitation and Proteomics Analyses

Spin-GFP protein was pulled down and interaction partners identified using a proteomics-MS approach; see supplemental section for schematic on workflow. Briefly, using the Gal4-UAS system, we overexpressed Spin-GFP under the control of the endogenous *spin* promoter (*spin*-Gal4; [20]. As a control we also used flies that had GFP being overexpressed with *spin-Gal4*. Using lysates from whole organisms, we immunoprecipated GFP using anti-GFP as the bait (GFP-Trap_A kit; Chromotek, Germany). The co-immunoprecipitated fractions were run on SDS-PAGE and each lane was cut into 10 slices, followed by in-gel digestion [50]. For each lane, slices were analyzed using 1200 nano-LC coupled to LTQ Orbitrap Discovery (Thermo Fisher Scientific, Germany) using Nanomate Triversa as ionsource. Peptide Identification and Protein assignments were performed using Protein Discoverer 1.3. (Thermo Fisher Scientific, Germany) and Mascot as search engine. Only proteins with a minimum of two unique peptides and false discovery rate better than 0.01 were accepted as positive hits. Proteins specific to Spin-GFP co-IP fraction and not found in the control-GFP co-IP fraction were compiled from three independent experiments.

#### Climbing Assays

Standard fly climbing assays were conducted to assay the climbing behavior in *spin* and control flies. For this, male and female flies were collected within 6 hours of eclosion and separately maintained in groups of 10-20 flies with transfers into fresh food every 2-3 days. At 3 days of age, these flies were subjected to a climbing assay and subsequently assayed for the next 30 days. Climbing assay was conducted in a double-blind procedure (for the entire procedure until the data analysis was completed) in a room devoid of any obvious olfactory and visual cues and temperature was monitored and maintained at 25**°**C. Briefly, flies were transferred to a glass vial with an 8cm mark on it. After acclimatization, the assay was video recorded starting with the tapping of the vial. Climbers were defined as flies that reached a height of 8cm within 18 seconds. The assay was repeated three times and averages for percent climbers was calculated from video recordings from a set of 3 independent vials/repeats with 10–20 flies each.

#### Life span analyses

Adult males and females of relevant genotypes were separately collected within 6 hours of emergence and kept in groups of 15-20. Flies were transferred into fresh food vials every 2 days and total number of surviving flies were recorded. These observations were used to calculate lifespan curves and 50% survivorship (T-50); which is the length in days at which 50% of the flies remained alive [51].

#### CHC extraction and analyses

Samples for were prepared by incubating 5-8 flies of each genotype at room temperature for 20 minutes with 120 µL of hexane containing 10 µg/mL hexacosane as an internal standard. 100 µL of the extract was transferred into a fresh glass vial and allowed to evaporate at room temperature. Samples were stored at −20 °C. Three replicates were prepared for each genotype.

Analysis by gas chromatography mass spectrometry (GCMS) was performed on a QP2010 system (Shimadzu, Kyoto, Japan) equipped with a DB-5 column (*5*%-Phenyl-methylpolysiloxane column; 30 m length, 0.25 mm ID, 0.25 µm film thickness; Agilent). Ionization was achieved by electron ionization (EI) at 70 eV. One microliter of the sample was injected using a splitless injector. The helium flow was set at 1.9 mL/min. The column temperature program began at 50 °C, increased to 210 °C at a rate of 35 °C /min, then increased to 280 °C at a rate of 3 °C/min. A mass spectrometer was set to unit mass resolution and 3 scans/ sec, from *m/z* 37 to 700. Chromatograms and mass spectra were analysed using GCMSsolution software (Shimadzu). For total CHC levels, the area under each of the CHC peaks were summed and normalized to the area under the peak for the spiked hexacosane standard. Statistical analysis was performed using a one-way ANOVA test with post-hoc Holm-Sidak test.

## Acknowledgement

Electron microscopy was conducted in the EM facility at the University of York, and the authors thank Megan Stark (Univ of York, UK) & Aurelien Dupont (MPI-CBG) for expert advice on TEM workflow & help with ultra-sectioning. Fluorescent imaging was carried out on microscopes housed in the Central Image and Flow Facility at NCBS and in the Light Microscope Facility at MPI-CBG. Shimadzu Asia Pacific provided use of a GCMS system. We are grateful to the NCBS Kitchen staff for help with fly culture media preparation, Jean-Christoph Billeter (University of Groningen) for providing the oenocyte-Gal4 driver, Marko Brankatschk (MPI-CBG) for generous donation of the anti-Lpp antibody, and Rachel Kraut (Biotec, Dresden) for providing comments and helpful suggestions during the course of this work. SH acknowledges E.Knust (MPI-CBG) for hosting her and providing resources to complete the experiments described here.

## Competing Interests

The authors declare no competing interests

## Author Contributions

SH & DS conceived the study, SH, STS & DS designed the experiments, SH, AK, RJ, YNC, JYY, STS, DS were involved in the experiments and data analyses. SH and DS wrote the manuscript with input from STS, JYY, and SJH.

## Funding

The authors wish to recognize the following sources of funding for this research: D.S. is supported by a Wellcome Trust/DBT India Alliance senior fellowship and is a recipient of NCBS–Merck & Co International Investigator Award. Work in the Sweeney lab was supported by a BBSRC studentship to SJH and a project grant (BB/I012273/1). YNC and JYY were supported by a grant from the Singapore National Research Foundation (NRF2010-06).

**Table 1:** List of identified Spin interactors

| No. | Peptide ID<br>(Uniprot) | Protein Name | Associated<br>Fly gene ID |
| --- | --- | --- | --- |
| 1 | Q01604 | Phosphoglycerate kinase | CG3127 |
| 2 | Q9VEB1 | Malate Dehydrogenase2 | CG7998 |
| 3 | P15007 | Enolase | CG17654 |
| 4 | Q0E9E2 | Pyruvate Carboxylase | CG1516 |
| 5 | B7Z0E0 | Isocitrate Dehydrogenase [NADP] | CG7176 |
| 6 | Q9W4H6 | GM13002p/ Pyruvate dehydrogenase | CG7010 |
| 7 | Q7K VX1 | Lethal (1) G0334, isoform C/ Pyruvate dehydrogenase | CG7010 |
| 8 | Q7K5K3 | GH08474p/ pyruvate dehydrogenase (acetyl-transferring) activity | CG11876 |
| 9 | P41572 | 6-phosphogluconate dehydrogenase | CG3724 |
| 10 | P52029 | Glucose-6-Phosphate isomerase | CG8251 |
| 11 | P52034 | 6-phosphofructokinase | CG6001 |
| 12 | Q9VHN7 | CG8036, isoform B/transketolase | CG8036 |
| **13** | **Q9V496** | **Apolipophorin** | **CG11064** |
| 14 | P11997 | Larval serum protein 1 gamma chain | CG6821 |
| 15 | O18404 | 3-hydroxyacyl-CoA dehydrogenase type-2 | CG7113 |
| 16 | Q9VSA3 | Probable medium-chain specific acyl-CoA dehydrogenase, mitochondrial | CG12262 |
| 17 | P02844 | Vitellogenin-2 | CG2979 |
| 18 | Q94511 | NADH-ubiquinone oxidoreductase, 75kDa subunit | CG2286 |
| 19 | Q7KN94 | Walrus, isoform A/FAD binding | CG8996 |
| 20 | Q95U46 | GH07925p/ flavin adenine dinucleotide binding | CG3902 |
| 21 | Q95U38 | GH10480p/ succinate-CoA ligase (ADP-forming) activity | CG11963 |
| 22 | Q9VNW6 | GH12632p/Glutamate 5-kinase | CG7470 |
| 23 | Q9VYD9 | AT04676p/ L-alanine:2-oxoglutarate aminotransferase activity | CG1640 |
| 24 | Q9VLC5 | Aldehyde dehydrogenase | CG3752 |
| 25 | P54399 | Protein disulfide-isomerase | CG6988 |
| 26 | Q9VC18 | CG11089/ IMP cyclohydrolase activity | G11089 |
| 27 | Q9VFC7 | Myofilin, isoform B/Mf5 | CG6803 |
| 28 | A1ZA66 | Stretchin-Mlck, isoform A | CG18255 |
| 29 | O18640 | Guanine nucleotide-binding protein subunit beta-like protein/Rack1 | CG7111 |

